# Enhancing transcriptome expression quantification through accurate assignment of long RNA sequencing reads with TranSigner

**DOI:** 10.1101/2024.04.13.589356

**Authors:** Hyun Joo Ji, Mihaela Pertea

## Abstract

Recently developed long-read RNA sequencing technologies promise to provide a more accurate and comprehensive view of transcriptomes compared to short-read sequencers, primarily due to their capability to achieve full-length sequencing of transcripts. However, realizing this potential requires computational tools tailored to process long reads, which exhibit a higher error rate than short reads. Existing methods for assembling and quantifying long-read data often disagree on expressed transcripts and their abundance levels, leading researchers to lack confidence in the transcriptomes produced using this data. One approach to address the uncertainties in transcriptome assembly and quantification is by assigning the long reads to transcripts, enabling a more detailed characterization of transcript support at the read level. Here, we introduce TranSigner, a versatile tool that assigns long reads to any input transcriptome. TranSigner consists of three consecutive modules performing: read alignment to the given transcripts, computation of read-to-transcript compatibility based on alignment scores and positions, and execution of an expectation-maximization algorithm to probabilistically assign reads to transcripts and estimate transcript abundances. Using simulated data and experimental datasets from three well-studied organisms — *Homo sapiens*, *Arabidopsis thaliana,* and *Mus musculus* — we demonstrate that TranSigner achieves accurate read assignments, obtaining higher accuracy in transcript abundance estimation compared to existing tools.

## Background

Long-read RNA sequencing (RNA-seq) represents a remarkable advancement towards achieving full-length sequencing of transcripts, offering novel insights into transcriptomes previously characterized only with short reads. Short-read sequencing data has limitations in several applications such as transcript assembly, primarily due to its fragmented nature and inherent biases (e.g., GC content, amplification) that add noise to downstream analyses (Benjamini & Speed, 2012; Hansen et al., 2010; Li et al., 2009). Long-read sequencing technologies address these limitations by substantially increasing the read lengths, allowing each read to generally cover a full-length transcript, and employing strategies such as direct RNA sequencing to reduce biases. Consequently, long-read data can provide more comprehensive and accurate profiles of complex transcriptomes.

However, despite their potential, the full capabilities of long-read RNA-seq remain untapped due to the limited inventory of tools optimized for analyzing long-read data. Although tools such as FLAIR (Tang et al., 2020), Bambu (Chen et al., 2023), ESPRESSO (Gao et al., 2023), and StringTie2 (Kovaka et al., 2019) are designed to characterize transcriptomes by both identifying novel isoforms and quantifying transcripts using long-read RNA-seq data, their results often lack agreement (Chen et al., 2023; Gao et al., 2023; Pardo-Palacios et al., 2023; Tang et al., 2020).

One way to address uncertainties in transcriptome assemblies is by assigning specific long reads to transcripts. This allows for a more in-depth evaluation of the read-level support for transcripts, as opposed to relying on read counts only. Given read-to-transcript assignments, transcripts can be directly associated with a distribution of supporting read lengths, quality scores, alignment positions, and more. These expanded sets of features can be used to derive a more confident set of transcripts and improve the accuracy of transcript abundance estimates.

Few tools, including FLAIR and Bambu, track read-to-transcript assignments, but this functionality is integrated into more complex pipelines that also identify novel isoforms in addition to quantifying known transcripts. A standalone tool capable of performing read assignment and quantification on any input transcriptome can be paired with other methods focusing on transcriptome assembly and could therefore enable users to investigate any transcriptome of their choice. However, this need remains largely unmet, with only a few recent methods, namely NanoCount (Gleeson et al., 2021), attempting to address it by quantifying transcripts, yet still lacking the ability to assign specific reads to transcripts.

Here we introduce TranSigner, a novel transcript quantification-only method that accurately assigns long RNA-seq reads to any given transcriptome. TranSigner first maps reads onto the transcriptome using minimap2 (Li, 2018, 2021) and extracts specific features from the alignments, such as alignment scores or the 3’ and 5’ end read positions on a transcript. These features are then utilized to compute compatibility scores between read and transcript pairs, which indicate the likelihood of a read to originate from a specific transcript. TranSigner then employs an expectation-maximization (EM) algorithm to derive maximum likelihood (ML) estimates for both the read-to-transcript assignments and transcript abundances simultaneously. We show that by guiding the EM algorithm in the expectation step with precomputed compatibility scores, TranSigner generates high-confidence read-to-transcript mappings and improves transcript abundance estimates.

## Results

### Simulated data performance

We first compared TranSigner against an existing quantification-only tool, NanoCount (Gleeson et al., 2021). We benchmarked all three tools using five sets of simulated ONT reads: three sets of direct RNA reads and two sets of cDNA reads. The reads were simulated from protein-coding and long non-coding transcripts in the GRCh38 RefSeq annotation (release 110), and then each tool was provided with both the simulated reads as well as the full RefSeq annotation as the target transcriptome (see Methods for a full description of the simulated datasets). For simplicity, we will refer to the transcripts from which the reads were simulated as the origin transcripts. To estimate how accurately a tool assigns a read to its respective origin, we conducted both linear and nonlinear correlation analyses between the expected read counts and each tool’s estimates, using Pearson’s correlation coefficients (PCCs) between raw read counts and Spearman’s correlation coefficients (SCCs) between log-transformed read counts, respectively. A linear correlation analysis evaluates the ability of a tool to assign each read to a transcript, while a nonlinear correlation analysis assesses how well estimates capture monotonic trends in gene expression patterns.

In both analyses, we observed that TranSigner’s estimates had stronger correlations with the ground truth compared to NanoCount’s, as illustrated in Figure 1, which shows results from one dataset typical of all three simulated ONT direct RNA datasets (see Supplementary Table S3 for the SCC and PCC values on each read set). In both log-transformed (Figure 1A) and raw (Figure 1B) read count correlation scatter plots, TranSigner shows higher concentrations of dots near the diagonal. However, this feature is not as pronounced in the plots of NanoCount’s results; the accumulations of dots well below the diagonal in the case of NanoCount reveal the tool’s tendency to underestimate the read counts. On the simulated ONT direct RNA datasets, TranSigner’s average SCC and PCC values were 0.867 and 0.999, whereas NanoCount’s were 0.667 and 0.997. TranSigner also achieves higher correlations with the ground truth when applied to the simulated ONT cDNA datasets (see Supplementary Figure S1, Supplementary Tables S4).

**Figure 1.**
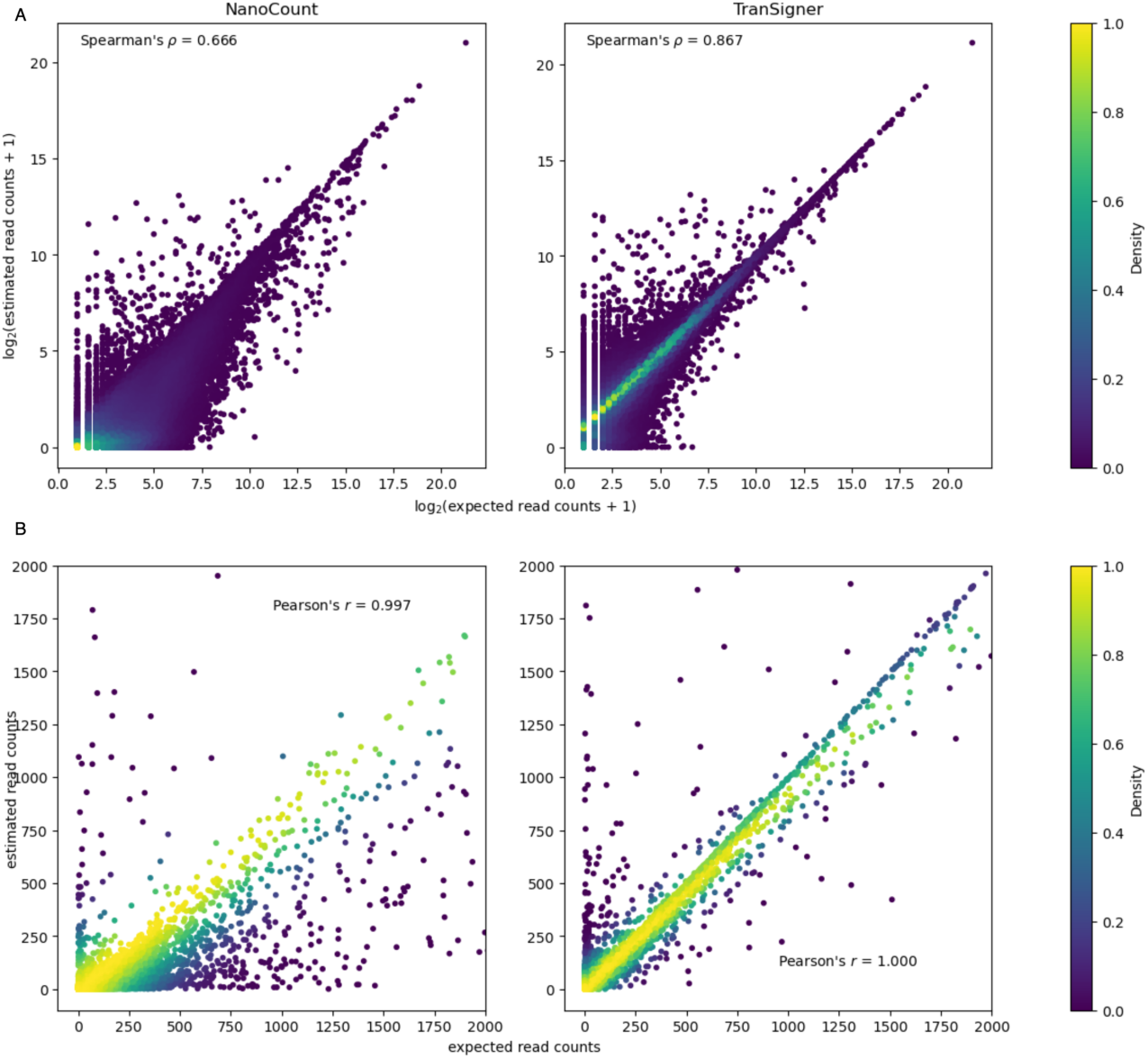
Correlation scatter plots comparing expected read counts to the read count estimates generated by NanoCount and Transigner on a simulated ONT direct RNA reads set. All tools were provided with the full RefSeq annotation from which the reads were simulated from. A: scatter plots showing the nonlinear correlations between the log-transformed ground truth and the estimated read counts. B: scatter plots showing the linear correlations between the raw ground truth and estimated read counts. The x- and y-axes were limited to [0, 2000] for demonstration purposes.

Even for extensively studied species, gene annotation catalogs are often incomplete, missing both potential gene loci and many transcript isoforms (Amaral et al., 2023; Varabyou et al., 2023). This is one reason why most long-read processing tools identify which transcripts are present before quantification.

Identifying novel isoforms not present in the annotation, as well as determining which of the known mRNA variants are expressed can lead to better quantification of expressed transcripts. This is illustrated by our results in Figure 2, where we show that the average nonlinear correlation coefficients between estimated and true read counts improve for both TranSigner and NanoCount when just the origin transcripts are provided in the input instead of the full reference annotation (see Supplementary Tables S3 and S4 for SCC and PCC values across all simulated ONT direct RNA and cDNA data sets).

**Figure 2.**
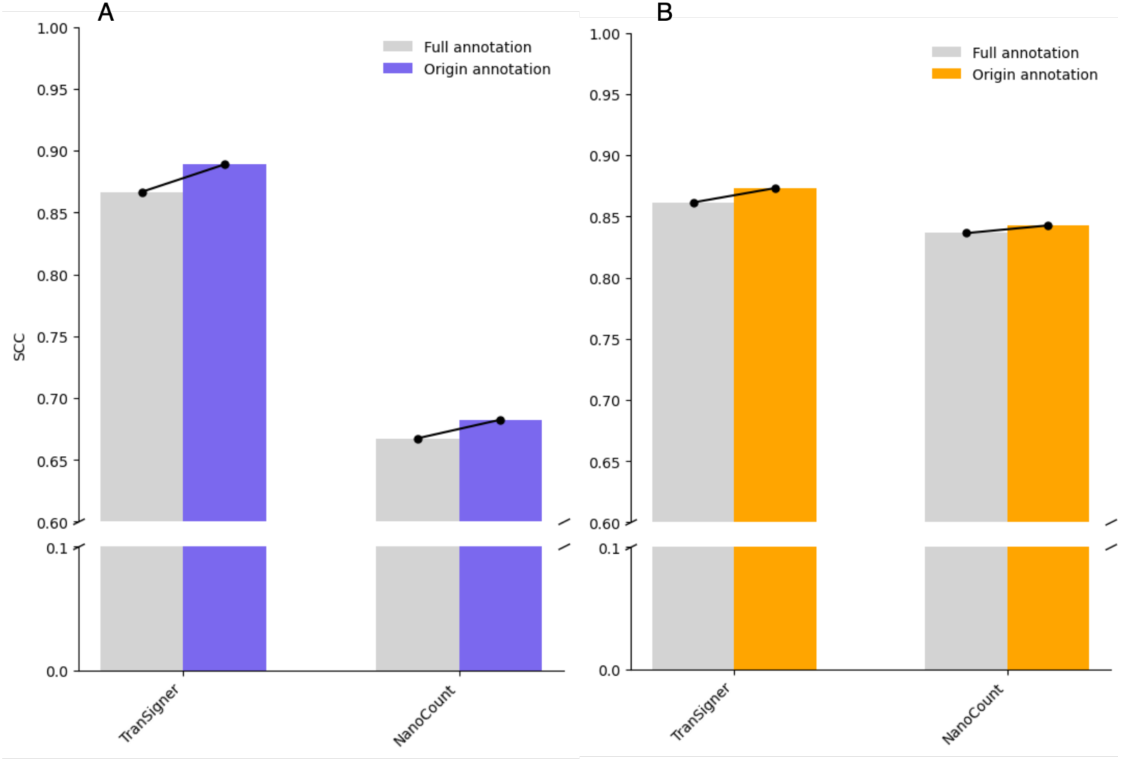
SCC values observed when either the origin transcriptome (blue in A, orange in B) or the full RefSeq annotation (grey) is used to run TranSigner and NanoCount on the simulated ONT reads. A shows the averages across 3 simulated ONT direct RNA read sets , while B shows the averages across 2 simulated ONT cDNA read sets.

Achieving an accurate transcriptome remains a challenging problem, with different tools obtaining varying accuracies in this task, while also relying to varying degrees on the input reference annotation. Using the same simulated ONT data sets (3 direct RNA, 2 cDNA) we used to benchmark TranSigner and NanoCount, we evaluated existing tools’ ability to handle incompleteness in the input guide annotations. To do this, we randomly sampled the full RefSeq annotation to include varying percentages–between 0% and 100% with increments of 5%–of the origin transcripts and provided the resulting annotations as guides to StringTie2, FLAIR, and Bambu. We did not include ESPRESSO in this comparison, as processing a single simulated data set took more than 24h to process. We also randomly sampled each percentage of retained origin transcripts three times (see Methods for further details).

Genome-guided transcriptome assemblers like StringTie2 (Kovaka et al., 2019) can reliably profile a transcriptome even in the absence of an input guide annotation, while methods like Bambu (Chen et al., 2023) or FLAIR (Tang et al., 2020) demonstrate a substantial decrease in both sensitivity and precision of transcript identification when the percentage of origin transcripts in the input guide annotation is progressively reduced. Figure 3 shows that while Bambu outperforms StringTie2 and FLAIR in terms of average sensitivity when a substantial portion of the origin transcriptome is provided in the input, StringTie2 consistently outperforms the rest of the tools in precision across all percentages of origin transcripts kept in the input annotation. Bambu achieved highest F1 scores when the guides retained most of origin transcripts, but StringTie2 gradually surpassed others as guides became increasingly incomplete (Figure 3C). Such resilience to varying degrees of incompleteness in the input transcriptome is critical, especially for studies involving poorly annotated organisms or in cases where the RNA-seq sample contains many novel isoforms (see Supplementary Tables S1 and S2 for the metric values on each dataset). However, StringTie2 does not assign individual reads to the transcripts it assembles, making it difficult for the user to check the reliability of the isoforms it assembles using long reads. By introducing TranSigner, we aimed to also address this gap, in addition to improving transcript quantification accuracies.

**Figure 3.**
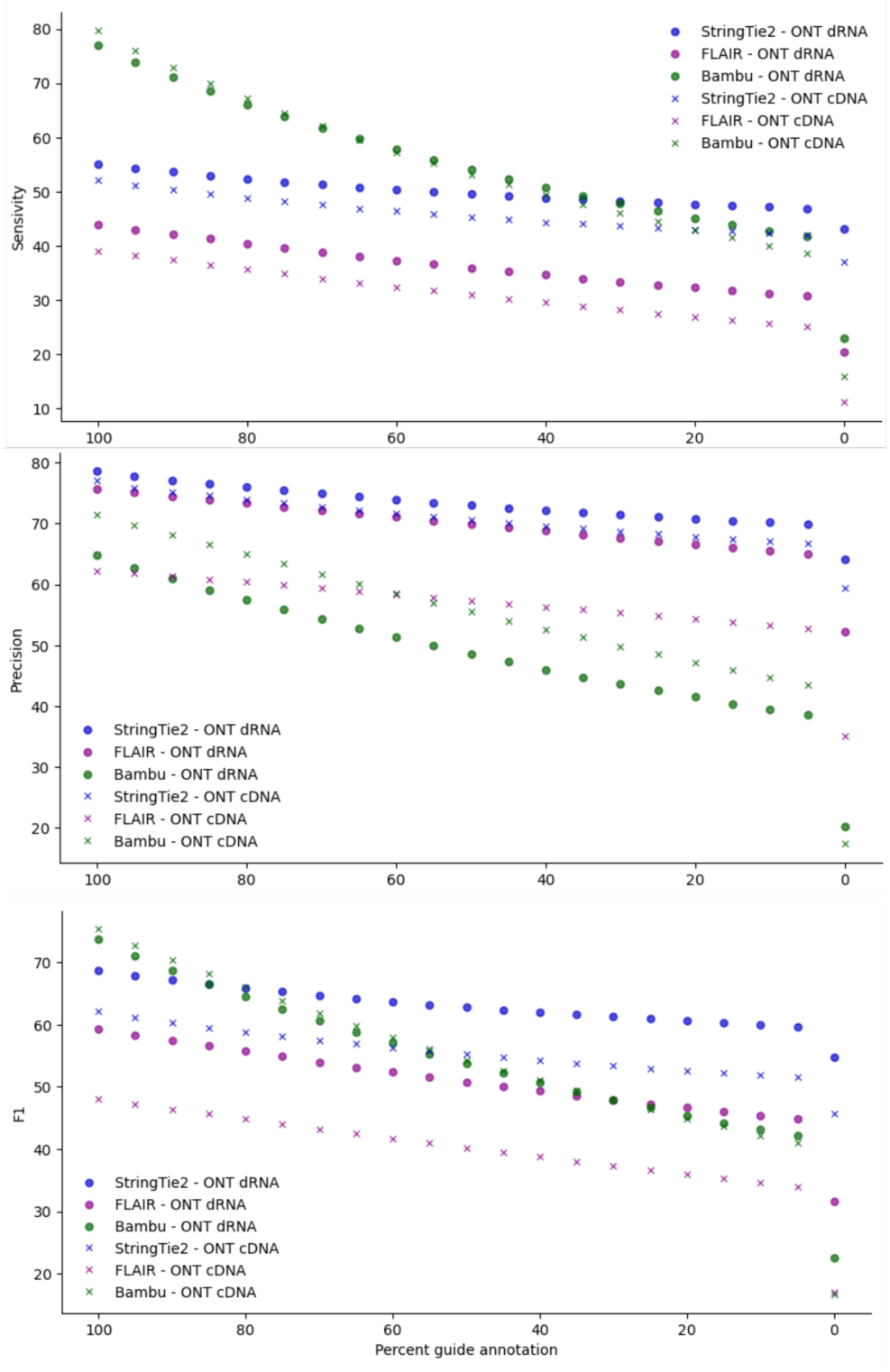
Long-read assembly accuracies of StringTie2, FLAIR, and Bambu with varying percentages (100% to 0%) of randomly sampled origin transcripts provided in the input guide annotation. Mean values for all three metrics – sensitivity, precision, and F1 – across ONT direct RNA and cDNA datasets are shown as circles and crosses, respectively.

Next, we compared TranSigner’s quantification accuracies against those of several other tools – StringTie2, NanoCount, Bambu, and FLAIR – when provided with guide annotations containing varying percentages of the origin transcripts. Since TranSigner is not capable of identifying novel transcripts, we also ran TranSigner on the transcriptome assembled by StringTie2 (denoted as StringTie2 + TranSigner) to investigate its performance against other tools, such as FLAIR or Bambu, which are capable of novel isoform identification. For this experiment, we re-used the same sets of simulated ONT reads and 5% ∼ 100% guide annotations sampled before.

Average correlation coefficients between the true and estimated read counts are shown in Figure 4 (also see Supplementary Tables S5 and S6 for results on all input datasets). Except for StringTie2 + TranSigner, every tool experienced a drastic drop in SCC values as the percentage of origin transcripts decreased. TranSigner had the highest correlation values when the input guide annotation contained nearly all origin transcripts. However, when 90% or fewer of the origin transcripts were retained in the guide annotation, StringTie2 + TranSigner yielded the best SCC values in both ONT direct RNA and cDNA benchmarks (Figure 4A, 4B), demonstrating that this combination is the best in preserving the rank of the expression values across most levels of incompleteness in the available annotation. This same pattern holds for PCC values (Supplementary Figure S2A, S2B). StringTie2 does not output read counts for its transcript abundance estimates, so it was excluded from this initial correlation analysis. As StringTie2 outputs read per base coverages, we post-processed TranSigner’s read-to-transcript assignments to generate read per base coverages (see Methods). TranSigner + StringTie2 obtains better read per base coverage PCCs (Figure 4C) and SCCs (Supplementary Figure S2C) correlation values than StringTie2. The improvement is more notable in PCCs than in SCCs.

**Figure 4.**
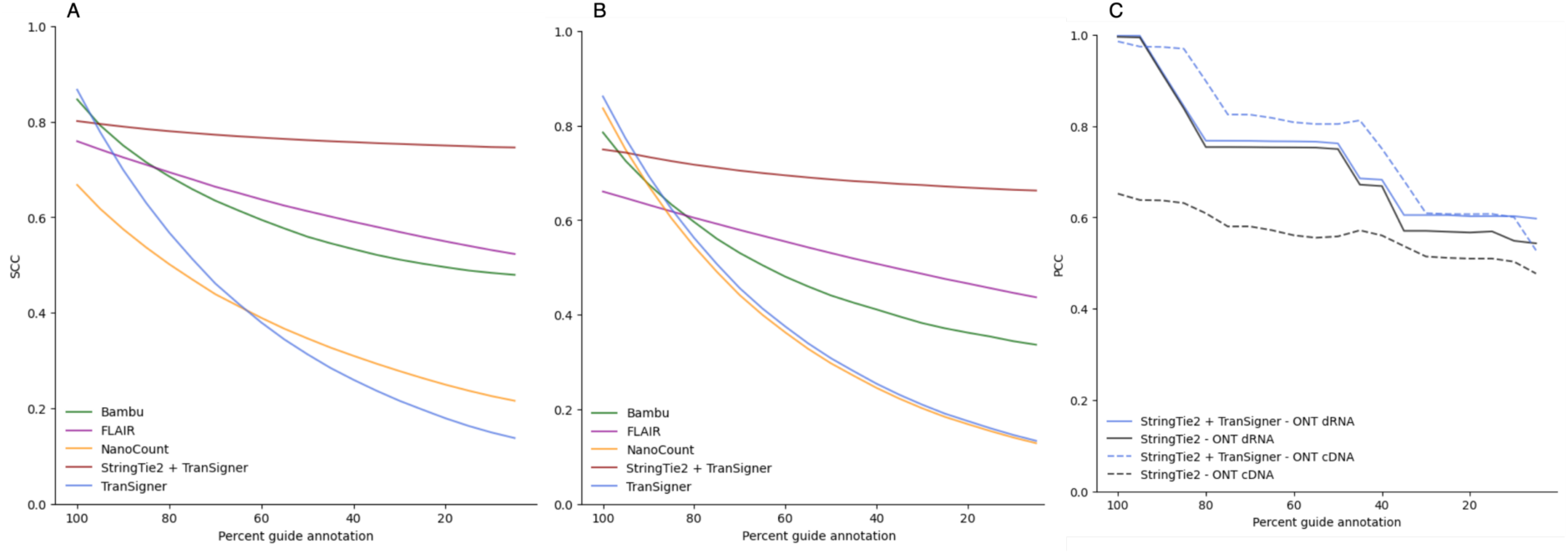
Correlation coefficients between true and estimated abundances (read counts in A and B, and per base read coverages in C) computed at varying percent guide annotations computed using simulated ONT data. A: SCC values in simulated ONT direct RNA data. Average SCCs across 9 independent observations (3 read sets, 3 guide samplings) shown. B: SCC values in simulated ONT cDNA data. Average SCCs across 6 independent observations (3 read sets, 2 guide samplings) shown. C: PCC values for both ONT direct RNA (solid line) and cDNA (dotted line) simulated reads. Averages across multiple samples are shown. Different colors indicate different tools.

One key feature of TranSigner is its ability to assign specific reads to transcripts, particularly useful in experiments where users need to identify reads originating from specific transcripts of interest. In this context, we compared TranSigner and StringTie2 + TranSigner with FLAIR and Bambu, which also output read-to-transcript assignments. Their performance was evaluated using recall, precision, and F1 scores, computed by counting the number of correctly versus incorrectly assigned reads (see Methods). When all origin transcripts are provided (i.e., 100% complete guide annotation), TranSigner demonstrated the highest sensitivity, recall, and hence F1 score (see Figure 5, Supplementary Tables S7 and S8). However, as soon as the guides become even slightly incomplete, StringTie2 + TranSigner had the highest performance, making it the preferred choice when the target transcriptome is 95% or less complete.

**Figure 5.**
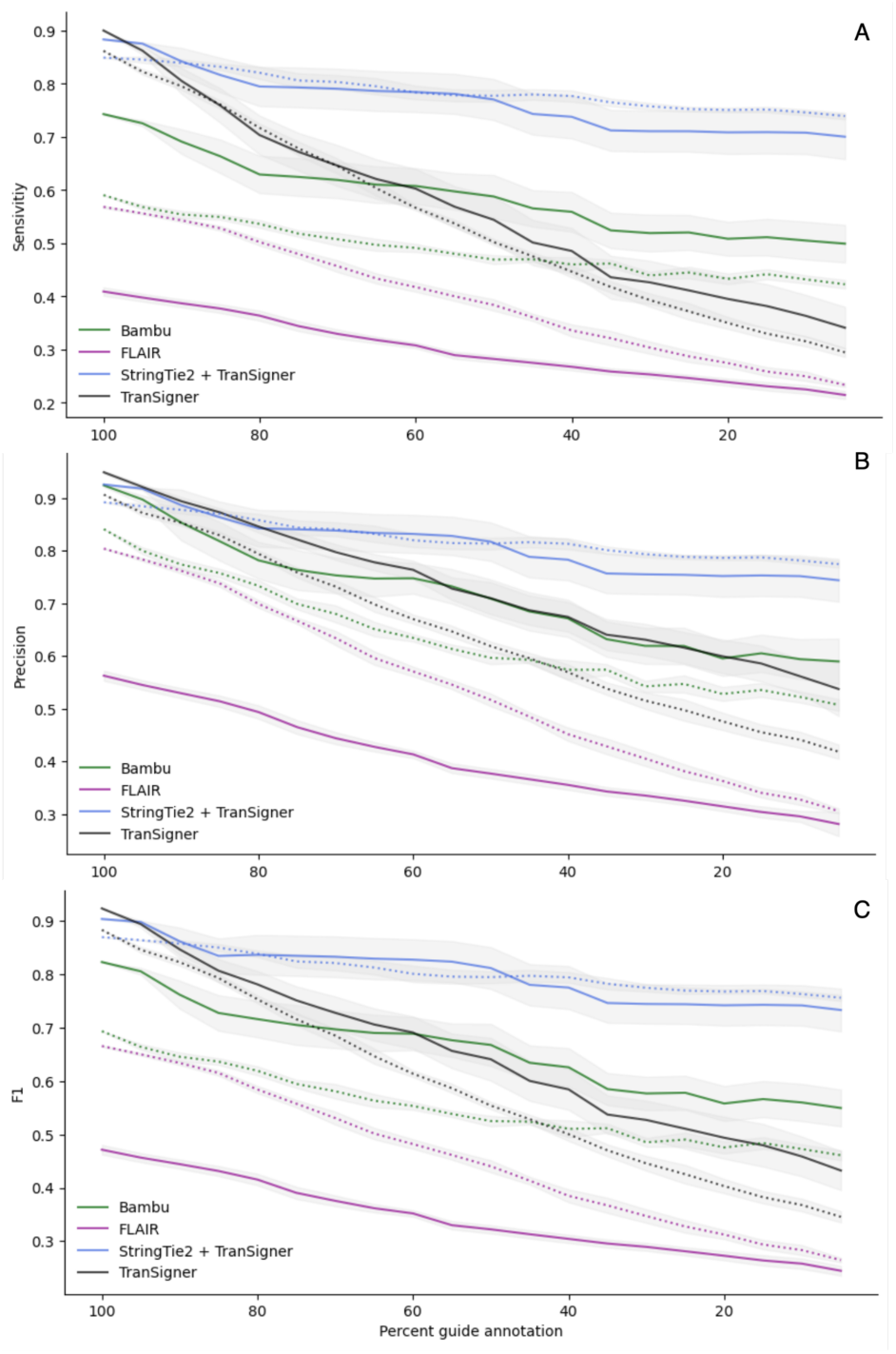
Read-to-transcript assignment accuracies for TranSigner, StringTie + TranSigner, Bambu, and FLAIR on simulated ONT data. Solid lines represent performance on ONT direct RNA reads and dotted lines represent performance on ONT cDNA reads. Three metrics – sensivitiy, precision, recall – are shown from top to bottom. Standard error of measurement (SEM) intervals are shown as shaded areas.

Although TranSigner achieved the highest F1 scores with nearly complete guides, its performance declined rapidly as the number of origin transcripts in the guides decreased, as expected (Figure 5C). A similar pattern of decline is observed in every tool across all metrics. Bambu experienced a greater drop in precision than StringTie2 + TranSigner, despite both starting at a similar value. Note that both Bambu and FLAIR showed fluctuations in performance depending on the ONT read types. In contrast, StringTie2 + TranSigner showed the least amount of variation in performance across different read types.

### Real data performance

To evaluate the performance of TranSigner and StringTie2 + TranSigner using experimental data, we utilized the ONT RNA-seq data sets provided by the Singapore Nanopore Expression Project (SG-NEx) (Chen et al., 2021), which include synthetic spike-in transcripts, known as sequins, with known annotation and concentrations. We selected 12 ONT direct RNA and cDNA samples from three different human cell lines: HCT116, K562, and MCF7. As ground truth for this experiment, we used the counts per million (CPM) values provided by SG-NEx and compared them with the estimates obtained by TranSigner, StringTie2 + TranSigner, and Bambu, the next best performer on the simulated data. We ran StringTie2 + TranSigner and Bambu twice, each time providing two different input guides: one including the full sequin annotation in addition to the GRCh38 reference annotation and the other containing only the GRCh38 reference transcripts without the sequins. The guide annotation without the sequins reflects real-world scenarios where transcript annotations are absent from the reference. TranSigner was only run with the full sequin annotation, as it cannot assemble any novel transcripts itself.

TranSigner achieved an average SCC of 0.88 between ground truth and estimated values, surpassing both Bambu (0.76) and StringTie2 + TranSigner (0.81) when provided with the full sequin annotation, as displayed in Figure 6 (also see Supplementary Tables S9). However, when no sequin annotation was provided, StringTie2 + TranSigner outperformed Bambu, obtaining an average SCC value of 0.64, compared to Bambu’s SCC value of 0.49. This trend persisted in linear correlation analyses, with TranSigner achieving the highest PCC value with full annotation (0.83), while StringTie2 + TranSigner’s was the best performer in the absence of sequin annotation (Figure 6 and Supplementary Tables S9).

**Figure 6.**
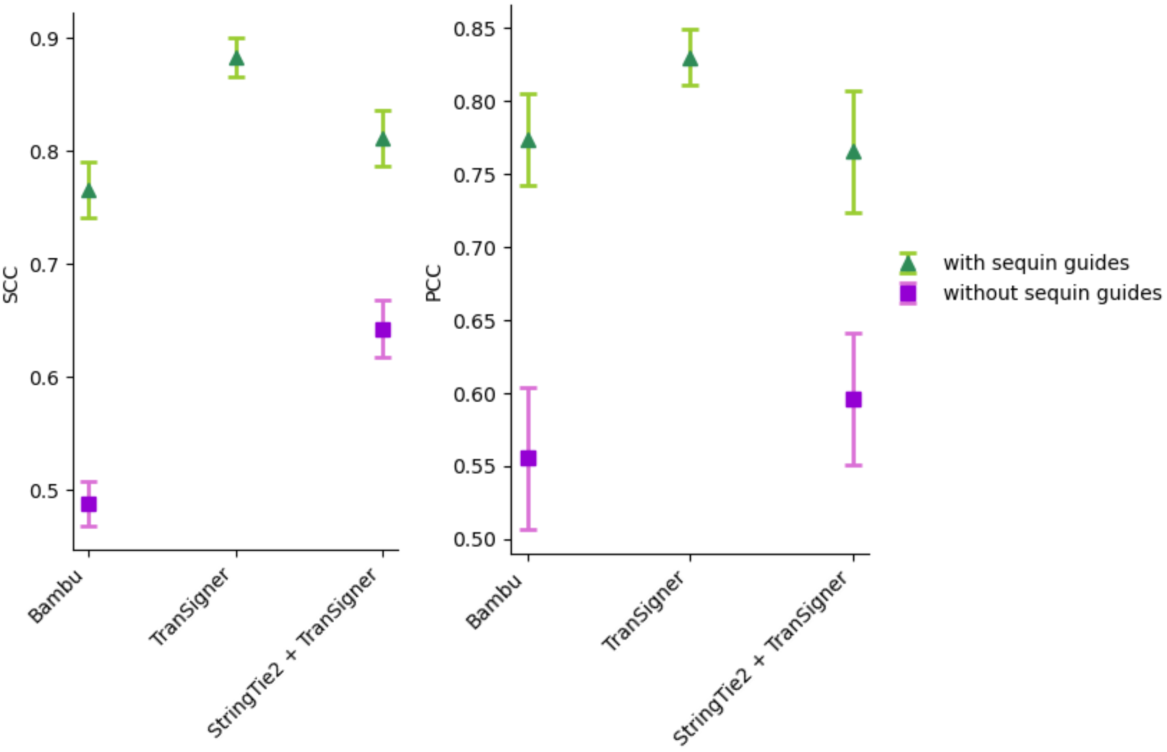
Correlation coefficients between estimated and expected sequin abundances measured using the SG-NEx data. Green triangles represent benchmark results when the full sequin annotation is provided, and purple squares when no sequins were present. SCC values are shown in the left, and PCCs on the right. Error bars represent the SEM values across 12 samples.

Overall, these results suggest that StringTie2 + TranSigner may be preferable in scenarios where numerous unannotated or novel isoforms are anticipated, while TranSigner is optimal when the reference is presumed to be nearly complete. Note that with complete sequin annotation, TranSigner outperformed both Bambu and StringTie2 + TranSigner, on all three different long-read types available in the data: direct RNA, direct cDNA, PCR-cDNA (average and per-sample SCC values shown in Supplementary Figure S3 and Supplementary Tables S9).

We also evaluated the correlation between short-read-based and long-read-based abundance estimates using publicly available paired short and long-read datasets, sequenced from the same biological sample. In all following results, the short-read libraries were all generated through poly-A selection and sequenced with Illumina sequencers, while the long reads were mostly generated using ONT direct RNA or cDNA sequencing protocols. Unlike the sequin samples or simulated long reads, the ground truth is unknown for these datasets as we lack information on which transcripts are expressed and their relative abundances. However, it is generally assumed that short reads provide more accurate abundance estimates compared to long reads, as they are less error-prone and typically yield more reads.

Specifically, we assessed the long read-based abundance estimates by two quantification-only tools we benchmarked with simulated data: NanoCount and TranSigner. All tools were provided with a StringTie2-assembled transcriptome, which represents a typical use for these tools where users provide transcriptomes assembled from samples of their interest. We used each tool’s abundance estimates to conduct nonlinear correlation analyses between the short read-derived TPM estimates and long read-derived CPM. As previously done for benchmarking long-read quantification tools (Pardo-Palacios et al., 2023), we assumed that a higher correlation between long read- and short read-derived abundance estimates is indicative of a higher quantification accuracy. Since none of the three quantification-only tools we used include TPMs in their output, we processed the read counts they provide to obtain counts per million (CPM) estimates, which are equivalent to TPMs in a long-read RNA-seq experiment where each read is considered to represent a transcript (see Methods for the read counts to CPM conversion equation). We used Salmon (Patro et al., 2017) to obtain TPM estimates on StringTie2 assemblies, using the Illumina short-read datasets (see Supplementary Text 3). As transcripts with low abundances are prone to misassembly and are often excluded from downstream analyses, we only included in our results transcripts with > 1 TPM as estimated by Salmon.

For our first experiment, we chose 21 short and long read paired datasets: 9 pairs from two normal human cell lines, A549 and HCT116, included in the SG-NEx datasets (Chen et al., 2021), and 12 pairs from two human cancer cell lines, H1975 and HCC827, provided by the long-read benchmarking of human lung cancer cell lines (Dong et al., 2023). The human lung cancer cell lines data sets also included PacBio reads, which are not present in the SG-NEx data sets. As shown in Figure 7, TranSigner consistently achieved higher correlations than NanoCount as well as StringTie2, across all read types (see Supplementary Tables S10 for the SCC and PCC values on each pair). TranSigner improved StringTie2’s estimates to varying degrees, with the highest improvements observed in the ONT PCR-cDNA data sets. Note that NanoCount was not evaluated on PacBio data as it was designed specifically to work with ONT data only.

**Figure 7.**
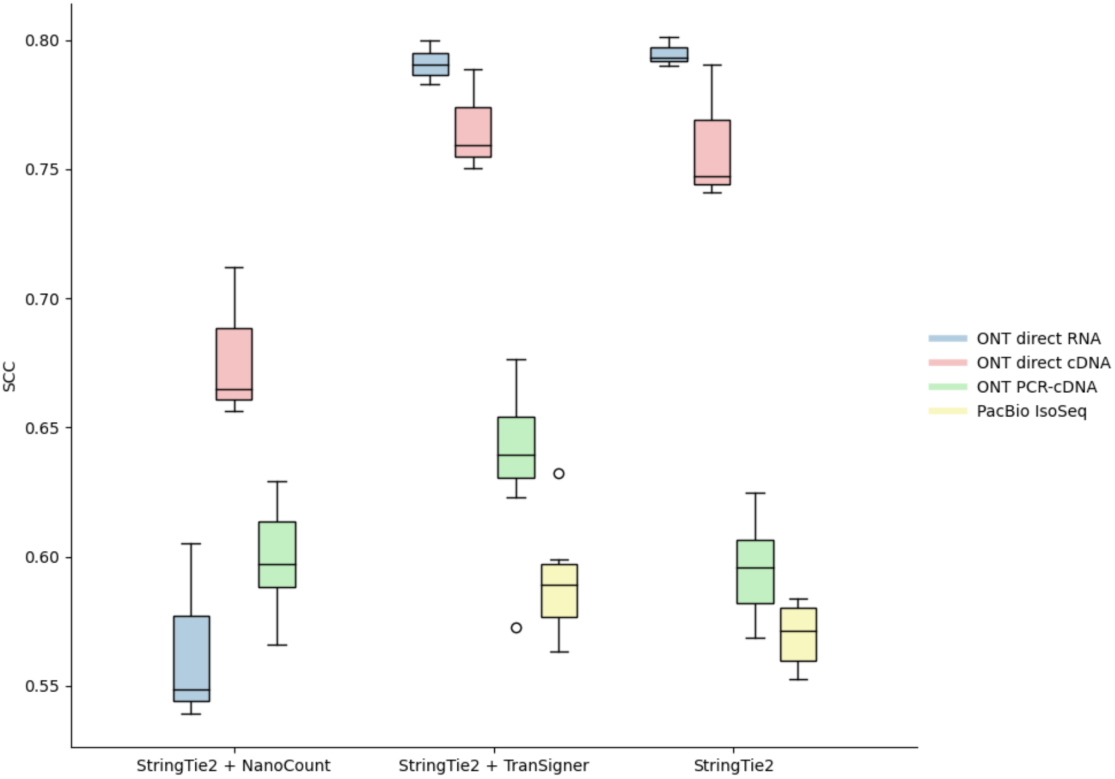
Box plots showing the distribution of SCC values between the short- and long-read-derived transcript abundances for 12 different pairs of human data sets. NanoCount and TranSigner were run on the StringTie2 assemblies on the long-read samples. StringTie2’s intial estimates are shown in the rightmost column for reference. Four distinct read types are shown in different colors.

Finally, we further expanded our benchmark to include paired short- and long-read data sets from two well-studies species: *A. thaliana* and *M. musculus*. To investigate how quantification accuracies vary at different levels of expression, we evaluated the performance of StringTie2 and StringTie2 + < a quantification-only tool > at progressively increasing TPM thresholds: 1, 5, 10, 15, and 20. For this experiment, we selected eight *M. musculus* pairs (four ONT direct RNA, four ONT cDNA) and three *A. athaliana* pairs (all ONT direct RNA). We benchmarked TranSigner’s and NanoCount’s performances when run on unguided StringTie2 assemblies, consistent with the previous analysis. As illustrated in Figure 8, when TranSigner was applied to StringTie2’s output, it achieved higher nonlinear correlations between short- and long-read TPM estimates than NanoCount, with the best improvements in SCC values obtained on the *M. musculus* ONT PCR-cDNA reads. These improvements were more pronounced for higher TPM thresholds.

**Figure 8.**
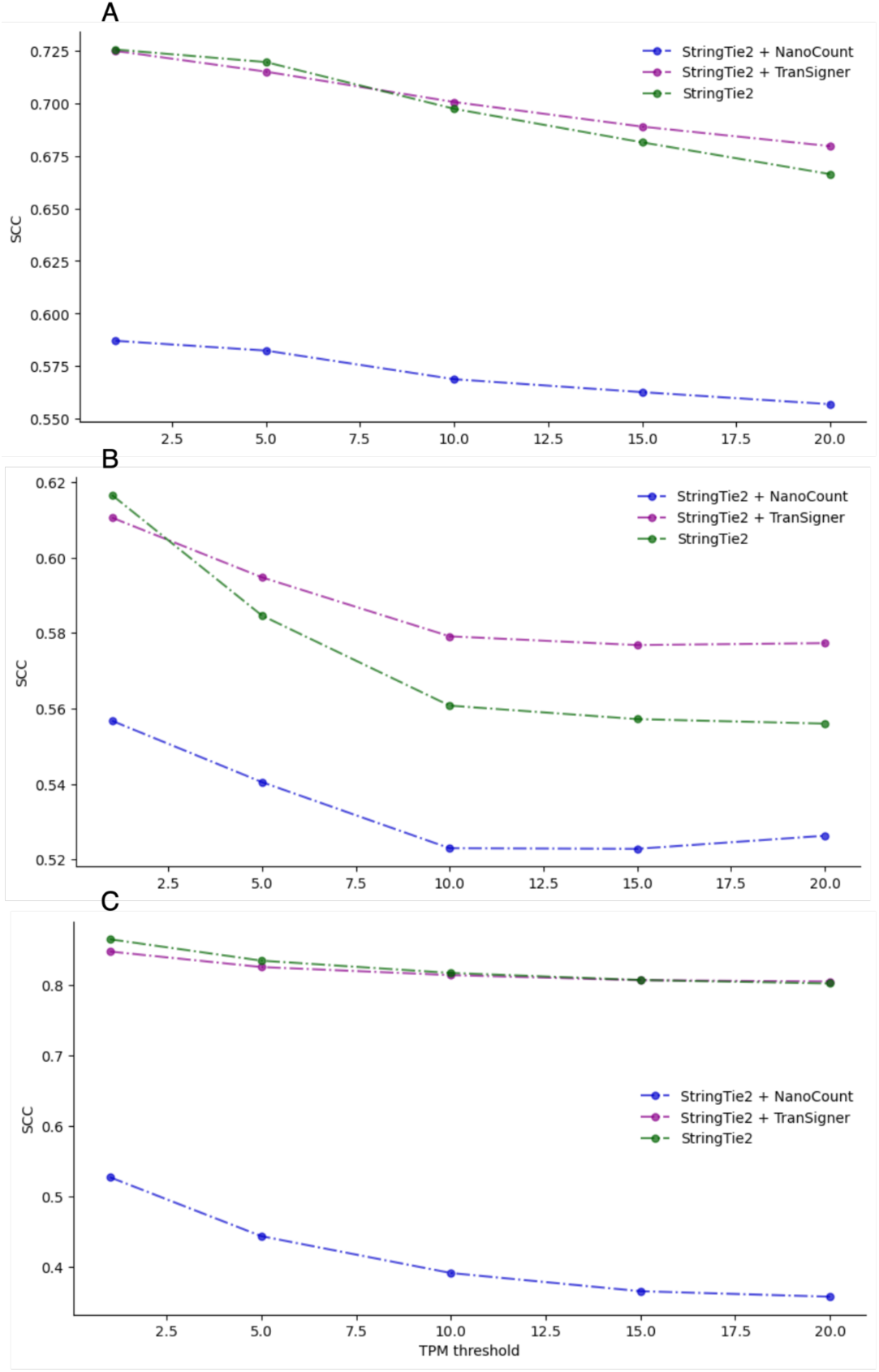
Correlation coefficient values between short- and long-read-derived transcript abundances estimated by NanoCount and TranSigner when run on StringTie2 assemblies, as well as StringTie2 itself, on paired *M. musculus* (A and B) and *A. thaliana* data sets (C). Each plot is showing a different organism and a different read type. A: average SCC values across increasing TPM thresholds on *M. musculus* ONT direct RNA data sets. B: average SCC values across increasing TPM thresholds on *M. musculus* ONT PCR-cDNA data sets. C: average SCC values across increasing TPM thresholds on *A. thaliana* ONT direct RNA data sets.

## Discussion and Conclusions

Assigning long reads to transcripts is a challenging task that involves the effective resolution of multi-mapping reads. Recent studies have unveiled the growing complexity of eukaryotic transcriptomes, revealing numerous isoforms across gene loci. The introduction of long-read RNA-seq technologies promises to uncover even more novel isoforms, as reads produced by these methodologies can capture full-length transcripts, overcoming the limitations of short reads. Although long reads cover transcripts at greater lengths, technical artifacts such as base calling errors and end truncations prevent these reads from being accurately mapped to their origins. With TranSigner, we have developed several strategies to address this challenge, facilitating the correct assignment of reads that ambiguously map to multiple isoforms.

Additionally, we designed TranSigner to complement another method capable of transcriptome assembly. As gene annotation is still an unresolved issue, determining the accuracy and completeness of a profiled transcriptome remains difficult. Users often struggle to select the appropriate reference for their analyses, leading to unpredictable impacts on their results. In our study, we observed a significant drop in assembly quality when less complete guides were provided. This suggests that tools heavily reliant on high-quality reference annotations may struggle in real-world scenarios where many novel isoforms are expected. By introducing a standalone tool for read-to-transcript assignments, we made these assignments easier to obtain regardless of the input transcriptome. Integrating this step into long-read RNA-seq data processing pipelines will improve the accuracy of transcriptomes identified using long reads by allowing users to inspect the quality of the reads supporting the transcripts and filter out less-supported transcripts. This, in turn, will lead to more accurate abundance estimates, as our results demonstrate the significant influence of assembly accuracy on correctly identifying transcript abundances.

## Methods

### Long-read RNA-seq model for read assignment

We describe the long-read RNA-seq process using a generative model (Figure 9). The conceptualization of RNA-seq as a generative process in which reads are sampled from a pool of transcripts has already been used in models for short-read quantification. We adopted the general framework proposed by others (Li et al., 2009; Pachter, 2011) but introduced necessary modifications to tailor the model to long read data. Given a read, we assume that three unobserved events in the RNA-seq experiment determine a read’s sequence: (1) the transcript from which that read was sequenced, (2) the position within the transcript of the 3’ end of the read, and (3) the transcript position of the reads’ 5’ end. Our model, thus, associates each observed read with three latent variables: the transcript (*T*) from which the read was generated, its 3’ end position (*S*), and 5’ end position (*E*) in *T*.

**Figure 9.**
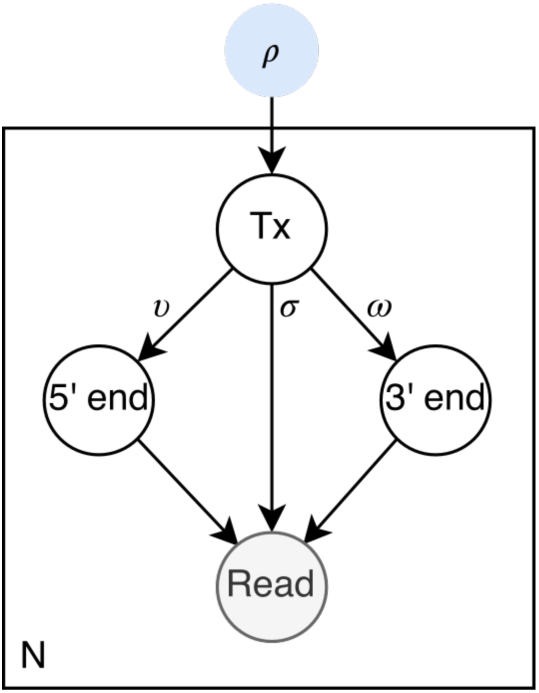
Graphical representation of TranSigner’s long-read RNA-seq model. Empty circles denote latent variables, the shaded circle represents the observed variable, and the blue circle indicates the primary parameter of the model – specifically, the relative abundance of the transcript. Parameters *υ*, *ω* approximate the likelihood of the specific 5’ and 3’ end positions of the read on the transcript, while parameter *σ* models the likelihood of observing a specific read sequence given a transcript and the read’s end positions. *N* represents the total number of reads generated in a single long-read RNA-seq experiment.

Existing RNA-seq quantification methods focus on accurately estimating *ρ*, the relative transcript abundances (Jousheghani & Patro, 2024; Li et al., 2009; Pachter, 2011). In contrast, our primary goal here is to assign reads to transcripts, which is solved by finding the most probable distributions over the latent variables, not *ρ*. However, deriving a maximum likelihood (ML) estimate on *ρ* also gets us ML estimates on the latent variable distributions, as they get repeatedly updated in the process of optimization. Hence, *ρ* is still the main parameter to optimize, and we define our objective with respect to *ρ* as follows. Given a set of transcripts *T* = {*t*} where |*T*| = *M*, the complete data likelihood function of our RNA-seq model is:

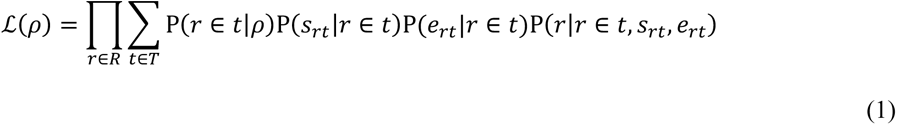

where *ρ* = {*ρ_t_*}_t∈T_ with ∑*_t_*_∈*T*_ *ρ_t_* = 1, *R* is the set of mapped reads defined as *R* = {*r*} with the cardinality of *N*, *s_rt_* and *e_rt_* are the 3’ and 5’ end positions of a read *r* in a transcript *t*, and *r* ∈ *t* indicates that *r* comes from *t*. Note Ρ(*O_r_* = *t*|*ρ*) = *ρ_t_*, since in an RNA-seq experiment the probability of selecting a transcript *t* to sequence depends on its relative abundance. We’ll approximate the 5’ end and 3’ end positions of a read in a transcript as the positions where the read alignment starts and ends on that transcript, respectively. The relationship between this likelihood function and read assignment estimates is easier to understand when Eq. 1 is rewritten as:

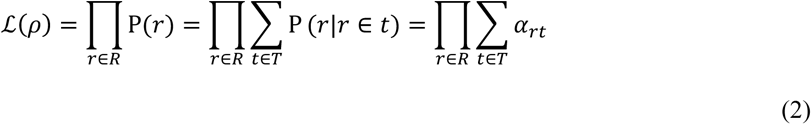

where *α_rt_* is the relative fraction of read *r* assigned to transcript *t*. Ρ(*r*) can also be written as a sum of conditional probabilities Ρ(*r*|*r* ∈ *t*), which represents the likelihood of *r* given that it comes from *t*. This conditional probability is also easily interpretable as the fraction of *r* that ought to be assigned to *t*, implying that a lower Ρ(*r*|*r* ∈ *t*) corresponds to a smaller *α_rt_*. Moreover, optimizing *L* involves driving *P*(*r*) to the maximum possible value in a probability distribution – 1, which is also equal to the sum of relative fractions of a read’s assignments to the set of transcripts (i.e., ∑*_t∈T_ α_rt_* = 1).

Different long-read RNA-seq technologies show various biases towards the ends of the transcripts (Amarasinghe et al., 2020; Chen et al., 2021; Grünberger et al., 2022; Wongsurawat et al., 2022). Nonetheless, long reads are more likely to cover all bases of a transcript, compared to short reads, which are generated from fragments of the transcript. The likelihood of a read’s end position should decrease as its distance from the transcript end increases. We model this expectation using two indicator variables– *υ* and *ω* for the 3’ and 5’ ends, respectively – to control how far apart the ends of a read can be from the ends of a transcript. For an alignment between a read *r* and a transcript *t*, we will refer to the distances between the alignment ends and transcript ends as ‘end distances’ and denote them as 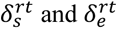 for the 5’ and 3’ ends, respectively. Then we define *υ* and *ω* as:

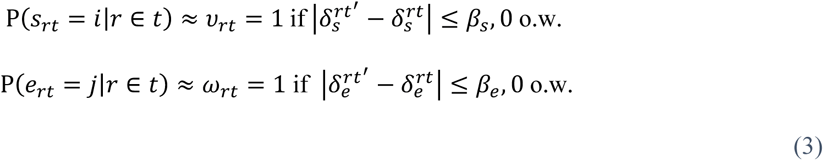

where 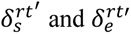 represent the end distances for the primary alignment of read *r* and transcript *t*′.

Here, *t*′ represents the transcript to which read *r* aligns on the primary alignment, which might not be the same as transcript *t*. Since alignment positions are indexed from the 5’ to 3’ direction on transcript *t*, end distances are computed as 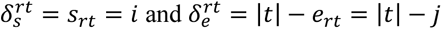 where |*t*| is the length of transcript *t*. Parameter *β* represents the tolerance threshold on how much greater the end distances can be compared to the primary alignment’s end distances for a given read *r*. This relative thresholding on end distances (*δ*) ensures that each read is compatible with at least one transcript (i.e., *t*′) after this filtering step since the primary alignment will always be considered “good,” which would not be true if a constant threshold was uniformly applied for all reads. When either *υ* or *ω* is set to 0, Ρ(*r*|*r* ∈ *t*) in Eq. 2 is also set to 0, and no fraction of *r* is assigned to *t*, guaranteeing that the corresponding (*r*, *t*) pair will be considered entirely incompatible, filtering it out from any downstream analysis.

Moreover, the parameters for the 3’ end are treated separately from those for the 5’ end because sequencing behaves differently at these ends. For example, there is a stronger coverage bias towards the 3’ end when nanopore-based direct RNA sequencing protocols are employed (Amarasinghe et al., 2020; Chen et al., 2021; Grünberger et al., 2022; Wongsurawat et al., 2022). We set the *β* parameter values based on both prior knowledge and a grid search (Supplementary Text 1). For the ONT direct RNA data, the current default values are *υ* = −∞ (i.e., no filter) and *ω* = −800, while for ONT cDNA and PacBio data, they are *υ* = −500 (i.e., unset) and *ω* = −550 for ONT cDNA and PacBio data.

The probability of observing a read *r* given all the latent variables is modeled using the alignment score between read *r* and transcript *t* (denoted by *x_rt_*) as:

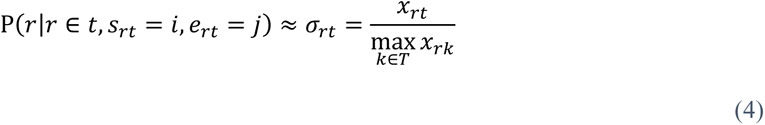

Note that if multiple alignments exist between read *r* and transcript *t*, we only retain the alignment with the maximum score. Using the above definitions, we can redefine the likelihood function as:

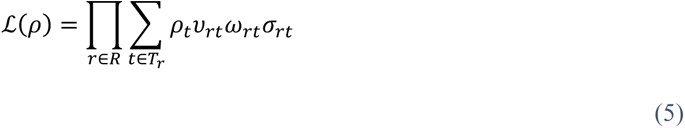

where *T_r_* is the set of transcripts aligned to read *r*, with *υ_rt_*, *ω_rt_*, and *σ_rt_* set to zero for any unaligned pair of read *r* and transcript *t*. By combining Eqs 2 and 5 we obtain that:

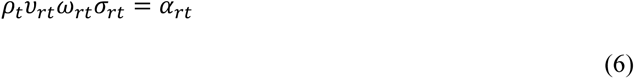

which shows how *α_rt_* can be computed from the alignments between reads and transcripts, assuming that the relative transcript abundances are given.

#### Alignment

We used minimap2 with parameter -N 181 to align the long reads to the set of input transcripts (Li, 2018, 2021). By default, minimap2 limits the maximum number of secondary alignments to 5. We observed that the number of true positives (correct read to transcript alignments) increases when we retain more secondary alignments, so we set -N to 181, the highest number of transcripts in a single gene locus according to the RefSeq release 110 annotation on the human GRCh38 genome, assuming this is the maximum number of secondary alignments a read can have. This strategy provides rough, preliminary estimates on the compatibility between reads and transcripts, without excluding any read and transcript pair for having suboptimal alignment scores. The user can freely adjust this parameter by specifying it in TranSigner’s input, which will then pass it to minimap2.

#### Alignment-guided expectation-maximization algorithm (AG-EM)

Our primary goal is to accurately assign reads to their respective transcript origins. We previously introduced *α* as a variable representing read-to-transcript assignments and established that the distribution over *α* is equivalent to that over the latent variables of our long-read RNA-seq model (Figure 9 and Eqs. 1, 2, 3). An expectation-maximum (EM) algorithm finds a maximum likelihood (ML) estimate for a main parameter (e.g., *ρ*) through iterative updates to the distribution over a set of latent variables (e.g., *α*). Hence, TranSigner employs an EM algorithm to obtain the most probable–in the sense that the complete data likelihood is maximized– distribution over *α* and presents the corresponding expected values as read-to-transcript assignments. It also outputs the ML estimates on *ρ*.

*Update rules*. The EM algorithm consists of alternating expectation (E) and maximization (M) steps, repeated until convergence. During the E step, the expected values for 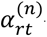–at some iteration *n*–are computed as follows:

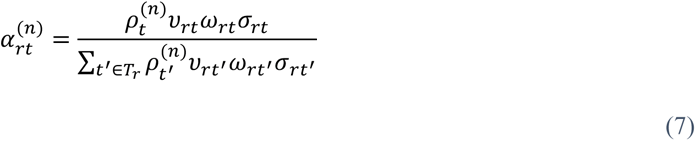

where *α* = {*α_rt_*}*_r_*_, *t*∈._ and *A* is the set of alignments between all reads and transcripts. In the following M step, then, the fragments of reads assigned to each transcript are summed up and then normalized by the total number of transcripts to get the relative transcript abundances, expressed as:

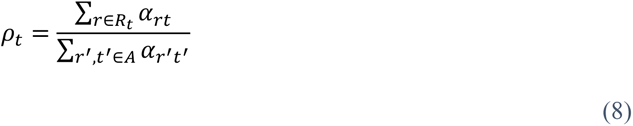

where *R_t_* is the set of reads aligned to transcript *t*. The denominator is constant across iterations and is equivalent to the total number of reads in a long-read RNA-seq experiment where each read represents a transcript, so we precompute this value before EM.

##### Initialization

Before the EM iterations, the relative transcript abundances (*ρ*) are initialized to the uniform distribution:

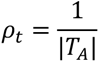

where *T_A_* is the set of transcripts with at least one alignment to a read in *R*. Additionally, the values for *υ*, *ω*, and *σ* don’t change during iterations, so we precompute their values and store them separately in a matrix *X* of dimensions *N* rows and *M* columns. For simplicity, we’ll refer to *X* as the compatibility score matrix. The computation specified in Eq. 7 is further simplified as:

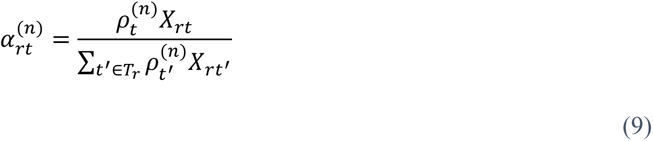

The pre-computation step involves a single scan over the alignment results, extracting values such as the alignment scores and alignment start/end positions, and then applying the definitions provided in Eqs. 3 and 4.

##### Optimization

Once *X* is precomputed and *ρ* is initialized, EM iterations are repeated until convergence, i.e., until the total sum of changes in the relative transcript abundances is less than a predefined threshold, by default set at 0.005. The user can adjust this threshold to increase the accuracy of the ML estimates at the expense of speed.

The novelty of our method comes from guiding the EM algorithm with the priors extracted from the alignment results, as detailed in the E-step update rule shown in Eq. 9. To further amplify the impact of these priors, we implemented an algorithm called the `drop`. The `drop` algorithm (Supplementary Figure S4) sets *X_rt_* = 0 if the fraction of read *r* that is assigned to transcript *t* (i.e., *α_rt_*) gets below a threshold, *τ* ∈ [0,1]. This effectively drops the compatibility relationship between read *r* and transcript *t* and ensures that no fraction of *r* gets assigned to *t* in any iterations following the drop, as *α_rt_* will always be 0 since its computation involves multiplication by *X_rt_* (Eq. 9). After the drop, another E-step is performed with the updated *X* scores to recompute the new *α_rt_* values. The *τ* value depends on the read *r* considered, and by default:

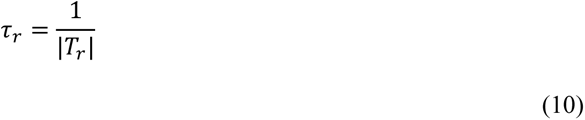

where *T_r_* is the set of transcripts that are compatible with *r*. The `drop` algorithm is called only right after the first E-step calculation, and its purpose is to discard minimap2 alignments that are not robust. The drop algorithm offers the potential to achieve a higher optimum compared to a naïve EM algorithm (Pachter, 2011), which relies solely on the relative transcript abundances (*ρ*) in its E-step update. We also allow users to increase this threshold (i.e., make it stricter) using the `-f` parameter that’ll increment *τ_r_* by a fraction of its own value as follows:

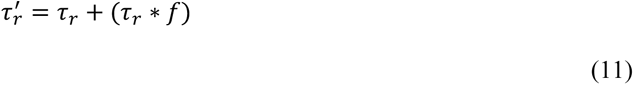

where *f* is a fractional value within the range [0, 1].

#### Read assignment

We can use the *α* values estimated by the EM algorithm to infer read assignments to transcripts. Raw *α* values represent fractional read assignments, where a single read may be distributed among multiple transcripts. These assignments might be challenging to interpret, as we assume each read to originate from a single transcript. To increase the interpretability and usability of the *α* values, we implemented the push algorithm (Supplementary Figure S5). This algorithm processes raw *α* values, converting them into hard assignments where each read is assigned to exactly one transcript. The push algorithm iterates through the reads and pairs each of them to the transcript with the highest read fraction as shown by the corresponding *α* value. It then recomputes the relative transcript abundances based on these hard assignments. These new *α* and *ρ* values may deviate from their EM-derived ML estimates, potentially resulting in reduced accuracy. We tested this using simulated data and observed only negligible reductions in accuracy.

### Implementation

TranSigner requires two inputs: a GTF file containing a reference gene annotation of the target transcriptome and a FASTQ file containing long RNA-seq reads. The reference annotation can be obtained from public sources such as RefSeq (O’Leary et al., 2016), GENCODE (Frankish et al., 2019), or CHESS (Varabyou et al., 2023), or it can be derived from transcriptome assemblies produced by programs like StringTie2. The latter annotations have the advantage of including novel isoforms while restricting the annotated transcripts to only those found to be expressed in the analyzed sample.

As illustrated in Figure 10, TranSigner consists of three modules: align, prefilter, and em. In the align module, input long reads are aligned to the target transcriptome using minimap2. The resulting alignment file becomes the input for the next module. Next, in the prefilter module, TranSigner extracts features such as the 3’ and 5’ end alignment positions and the ms alignment scores computed by minimap2. These features are used to compute the compatibility score matrix between transcripts and reads, as well as an index of the IDs of the transcripts found to be compatible with reads in the align module, which represent a subset of the target transcriptome.

**Figure 10.**
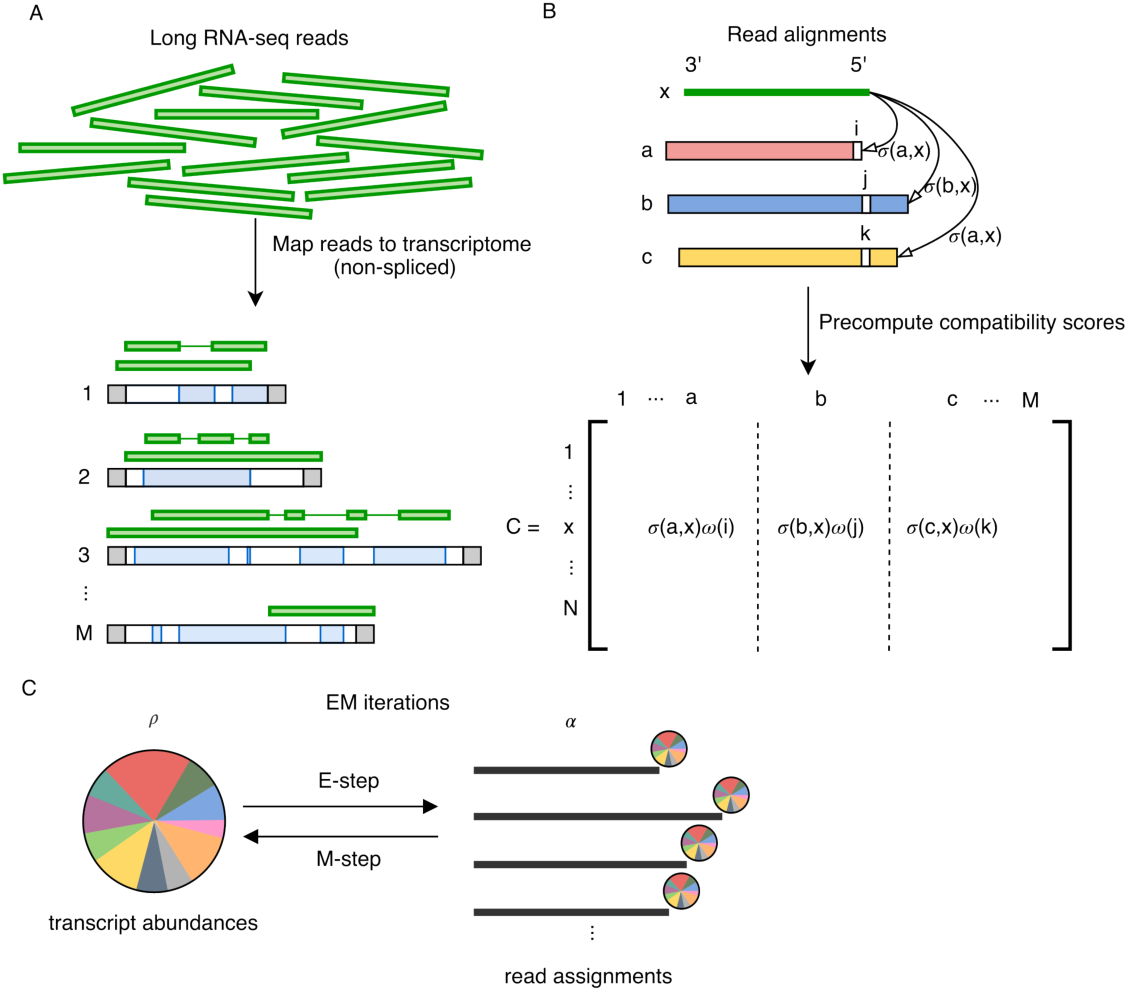
TranSigner’s workflow consists of three modules: align, prefilter, and EM. A: In the align module, N reads are mapped to M transcript sequences; B: In the prefilter module, compatibility scores are precomputed, and some alignments are filtered out; C: In the EM module, read fractions are assigned to transcripts and transcript abundances are updated iteratively until convergence.

Finally, the EM module takes as inputs the compatibility score matrix and the target transcriptome index from the prefilter module. It estimates the transcript coverage abundances using an expectation-maximization (EM) algorithm. The EM algorithm converges when the total change in the relative transcript abundances (*ρ*) is less than a specified threshold, by default set to 0.05. The drop algorithm, described above and in Supplementary Figure S5, is implemented as a component of this module. It allows users to use the `--drop` flag to remove low compatibility relations between reads and transcripts immediately after the first E-step update. Read-to-transcript assignments (i.e., *α* estimates) and relative transcript abundances (i.e., *ρ* estimates) are outputted as TSV files at the end of the EM module. Users also have the option to further process the assignments and output hard 1-to-1 assignments between reads and transcripts for increased interpretability by specifying the `--push` flag, whose algorithm is described in Supplementary Figure S5.

### Simulated data

Three sets of Oxford Nanopore Technologies (ONT) direct RNA reads and two sets of ONT cDNA reads were simulated using NanoSim (Gleeson et al., 2021). Expression levels were derived from protein-coding and long non-coding transcripts located on the main chromosomes (i.e., chromosomes 1 – 22, X, and Y) of the GRCh38 genome, extracted from the RefSeq annotation (release 110). We supplied the NA12878 direct RNA and cDNA reads from Workman et al. to NanoSim’s read characterization module to first construct two separate read profiles, one for generating direct RNA and the other for generating cDNA reads (Workman et al., 2019). We then estimated the transcript abundances of the direct RNA and cDNA samples by aligning each sample to the GRCh38 genome using minimap2 and providing the alignment results to salmon (Patro et al., 2017) in its alignment-based mode. We used the RefSeq annotation as the target transcriptome. Salmon estimates were then used as input for the NanoSim simulation module. For each direct RNA read set, we generated ∼14 million ONT direct RNA reads, and ∼25 million for each cDNA read set (Supplementary Text 5).

### Spiked-in data

We used an ONT direct-RNA dataset, which was released as part of the Singapore Nanopore Expression Project (SG-NEx) (Chen et al., 2021). This dataset was sequenced from three different human cell lines, HCT116, K562, and MCF7, and includes synthetic sequencing spike-in RNAs, also known as sequin RNAs. We used the SG-NEx-provided genome, which includes the in silico chromosome on which sequins are defined, to align these datasets. We also obtained the sequin transcripts annotation, their raw abundances, and the sample-wise spike-in concentration (i.e., from the SG-NEx AWS repository). To obtain sequin counts per million (CPM) levels, we followed the same method as in Chen et al..The ground truth sequin CPM for a sequin transcript *x* in a given sample *s* was computed as follows:

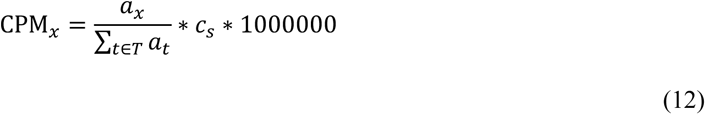

where *a* is the set of raw abundances provided by SG-Nex, *t* iterates through the entire set of transcripts to get the sum of all abundances, and *c_s_* is the spike-in concentration in sample *s*.

### Paired short- and long-read RNA-seq data

For humans, we employed paired short- and long-read RNA-seq data from the SG-NEx collection and long-read transcriptome profiling of human lung cancer cell lines data sets. Short- and long-read datasets are considered paired if they were obtained by sequencing the same biological sample. A subset of these samples included spike-in RNAs, and their reads were aligned to augmented versions of the GRCh38 genome that also includes the sequin-containing in silico chromosomes, provided by the original authors. All other samples (i.e., not spiked) were aligned to the regular GRCh38 p13 genome.

The goal with paired RNA-seq data sets is to compute the correlation between the short- and long-read-derived transcript abundance estimates. Long reads are first aligned to the GRCh38 genome using minimap2 and the resulting alignments are provided to StringTie2 for a transcriptome assembly. Short reads are then quantified on the long-read-derived StringTie2 transcripts using Salmon. Afterward, we ran quantification-only methods – NanoCount and TranSigner – on the StringTie2 assembly to obtain long-read-derived abundance estimates. We evaluated these tools’ estimates based on their nonlinear correlation with Salmon’s short-read-derived estimates (see Supplementary Text 3 for the commands used for short-read quantification). We repeated the same steps for two other organisms: *A. thaliana* and *M. musculus*. None of the samples from these two species contained sequins, so all reads were aligned to their respective reference genomes.

### Read assignments evaluation

For simulated and sequin data, we can define the following values based on the known origin transcript of each read:

- True positive (TP): a read is correctly assigned to its true origin.
- False positive (FP): a read is incorrectly assigned to a transcript that is not its true origin.
- False negative (FN): a read is not assigned to its true origin.

If a read is assigned to multiple transcripts without specifying the fraction allocated to each transcript, then the read is evenly distributed among those transcripts, with these fractions contributing to TP and FP values as appropriate. If the exact fraction of a read assigned to a transcript is provided, those fractions are used instead.

For each sample, the recall value of a method for the read-to-transcript assignment is calculated as the number of TPs divided by the total number of reads sequenced from that sample. The precision value is computed as the number of TPs divided by the sum of TPs and FPs. F1 score is defined as 2 * precision * recall / (precision + recall).

### Transcript abundance estimates evaluation

By default, TranSigner outputs read counts and relative transcript abundances as its quantification estimates. The read count of a transcript *t* (denoted as rc_t_) is the sum of all positive read fractions assigned to transcript *t*, while the relative transcript abundance of *t* (denoted as *ρ*_t_) is equal to rc_t_ normalized by the sum of all transcript read counts, ensuring that ∑_t∈T_ *ρ*_t_ = 1. Note that in a long-read RNA-seq experiment, each read counts as a transcript, making the sum of the read counts equivalent to the total number of transcripts identified from the long-read data.

TranSigner’s read count estimates can be converted to counts per million (CPM) estimates by calculating CPM_t_ = rc_t_/*l* * 10^6^ where *t* is a transcript and *l* is the total number of reads (aligned and unaligned). TranSigner also outputs read-to-transcript assignments where each read is assigned to one or more transcripts. More precisely, TranSigner outputs a list of transcripts to which a read *r* is assigned along with the fraction of *r* assigned to each transcript in that list, or the *α* estimates. These assignments can be used to compute coverage estimates for transcripts as 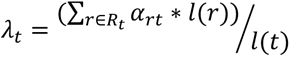 where *α_rt_* is the fraction of *r* assigned to transcript *t*, *R* _t_ is the set of reads whose fractions were assigned to *t*, and *l* is a function that returns the length of a read or a transcript.

We performed both linear and nonlinear correlation analyses to evaluate the correlation between estimated and ground truth values, each assessing different qualities of the read assignment and quantification methods. While nonlinear correlation analysis, utilizing log-transformed read counts and Spearman’s correlation coefficient (SCC), evaluates monotonic trends in the data, linear correlation analysis, utilizing Pearson’s correlation coefficient (PCC), assesses a tool’s accuracy in assigning all reads to transcripts, valuing each read equally regardless of its source. It’s worth noting that log transformation is typically applied to reduce variance in gene expression values. However, log transformation may compress differences in data points with large magnitudes, potentially diminishing the impact of errors in assigning reads to high abundance transcripts.

### Evaluation of tools capable of transcriptome assembly

We assessed the quality of assemblies generated by StringTie2, Bambu, and FLAIR using the intron chain-level sensitivity and precision values computed by GffCompare (Pertea & Pertea, 2020). We initially wanted to include ESPRESSO in this comparison, but we were unable to run it as it took more than 24 hours to process a single sample containing ∼14 million reads.

We benchmarked each tool using random samples of the RefSeq annotation to observe how well the completeness of the guides impacts the accuracy of the assembled transcriptome and the simulated ONT data. More precisely, we randomly sampled a percentage of the origin transcriptome, referring to the set of transcripts from which a set of reads are simulated, to remove from RefSeq. The guides were sampled to contain 21 different percentages between 0% and 100% of the origin transcriptome. For each percentage, we independently sampled the guides three times, yielding 63 different guides per read set. StringTie2, Bambu, and FLAIR were provided with the same guide annotations. Additionally, StringTie2 and Bambu were provided with the same minimap2 alignment results produced using the recommended options for processing ONT RNA-seq data (`-x splice -uf -k14` for direct RNA reads and `-x splice` for cDNA reads); FLAIR had its own align module. Unlike StringTie2 and FLAIR which output an annotation containing only the identified expressed transcripts, Bambu outputs both expressed and unexpressed transcripts in the guide annotation (see Supplementary Text 2). Therefore, for our evaluations, we removed any transcript that was assigned a zero read count from Bambu’s output.

## Declarations

### Ethics approval and consent to participate

Not applicable.

### Consent for publication

All authors have consented for publication.

### Availability of data and materials

The *A. thaliana* and *M. musculus* datasets are available from the European Nucleotide Archive (ENA) under accession numbers PRJEB32782 and PRJEB27590. Specific ENA sample accession IDs for each pair of short- and long-read data sets are made available in Supplementary Table S11. The SG-NEx samples containing spike-in RNAs are available from GitHub, ENA, and AWS open data registry. The long-read benchmarking on the human lung cancer cell lines data sets are made available from Gene Expression Omnibus (GEO) under accession number GSE172421. The values used to generate plots in this manuscript are made available as Supplementary Tables S1 ∼ S11. Supplementary Tables S0 contains the captions for each table. TranSigner is implemented in Python and is publicly available at https://github.com/haydenji0731/transigner and is also archived in Zenodo at https://doi.org/10.5281/zenodo.13334738. All code used to generate all figures (either in the main manuscript or in the supplementary materials) and the scripts and data files (e.g., ground truths for simulated and sequin data) used for benchmarking are available in Zenodo at https://doi.org/10.5281/zenodo.13334733. The transcript abundances used for read simulation are also available at the same address.

### Competing interests

The authors have declared no competing interests.

### Funding

This work was supported in part by the US National Institutes of Health under grants R01-HG006677, and R01-MH123567. The funders had no role in the study design, data collection and analysis, decision to publish, or preparation of the manuscript.

### Authors’ contributions

HJJ and MP designed the study. HJJ wrote the software and code used for benchmarking. HJJ and MP evaluate analysis results and wrote / revised the manuscript. HJJ prepared all (main and supplementary) figures. All authors read and approved the final manuscript.

## Acknowledgments

We would like to thank Jennifer J. Lee for proofreading this manuscript, Beril Erdogdu for engaging in discussions on long-read RNA-seq models, and Ales Varabyou for giving invaluable insight into experimental setups.

## Notes

### Summary of Updates

Figures 1, 2, 4, 7 and 8 updated; main text updated accordingly; supplementary tables and texts also updated.

