## Supplementary Texts and Figures for "Enhancing transcriptome expression quantification through accurate assignment of long RNA sequencing reads with TranSigner"

**1. Parameter tuning for 3' and 5' end distance thresholds.** TranSigner filters out the alignment between a read $r$ and a transcript $t$ if it starts and/or ends at a position far away from the 5' and 3' ends of $t$, indicating the aligned segment has a relatively small target cover. For simplicity, we introduced the concept of 5' and 3' end distances quantifying the number of unaligned base pairs at either end of $t$ (see Methods). To keep at least one alignment per read (i.e., its primary alignment), we threshold on the degree of increase in end distances, or $\delta$. For instance, if the 5' end distance for the primary alignment of $r$ is 120 bp. Then, an alignment with a 5' end distance of 150 would yield $\delta_s = -30$ where the subscript $s$ just indicates that the distance is at the start / 5' end of $t$.

Based on the prior knowledge of frequent 5' end truncations in ONT direct RNA reads, we set the 5' end threshold, $\beta_s$, to a lenient value of -800, and no 3' end filtering. Relatively little is known about the read distributions in ONT cDNA data, so we decided to tune the filter thresholds on a set of simulated ONT cDNA reads. We aimed to set the default values for $\beta_s$ and $\beta_e$ with ones that obtain the most accurate abundance estimates, as measured by their non-linear correlation with the known ground truth in the simulated ONT cDNA data. Briefly, we performed a grid search with $\beta_s$ values ranging from -550 to -600 with decrements by 50 and $\beta_e$ ranging from -300 to -500 with a decrement size of 100. We set $\beta_s$ to -600 and $\beta_e$ to -500 (see Supplementary Text Table 1).

| $\boldsymbol{\beta_s}$ **(-fp)** | $\boldsymbol{\beta_e}$ **(-tp)** | **Spearman's Correlation Coefficients (SCCs)** |
|---|---|---|
| -550 | -300 | 0.6908945215530291 |
| -600 | -300 | 0.6913623708965158 |

| | | |
|---|---|---|
| -550 | -400 | 0.6918556563135633 |
| -600 | -400 | 0.692299875863824 |
| -550 | -500 | 0.6923891562585597 |
| -600* | -500* | 0.6927918346209779 |

**Supplementary Text Table 1. SCCs observed in the $\beta_s$ and $\beta_e$ grid search on simulated ONT cDNA data.**

*TranSigner's current recommended -fp and -tp parameters when processing ONT cDNA or PacBio samples.

**2. minimap2 alignment parameters.** The long-read analysis tools we benchmarked require that long reads are aligned to either the genome or the transcriptome of interest. StringTie2, FLAIR, and Bambu use genome alignments, whereas NanoCount and TranSigner take in transcriptome alignments. FLAIR has its own align modules, so instead of running minimap2 ourselves, we ran this in-house module for benchmarking. For others (i.e., StringTie2 and Bambu), we used the minimap2-recommended parameters for aligning either ONT dRNA or cDNA reads to a genome (see Supplementary Text Table 2). As for transcriptomic alignments, NanoCount utilizes the minimap2 parameter preset for the alignment of genomic ONT reads but with an increased -N value. TranSigner uses an even higher -N parameter (i.e., 181) to retain all secondary alignments as described in Methods. 181 is the highest number of transcripts in a single gene locus according to the RefSeq release 110 annotation on the human GRCh38 genome. Users can adjust this parameter to an even higher number as needed, or provide a reference annotation to re-calculate the highest number of transcripts in a single gene locus. Since RNA-seq reads are being aligned to fully processed transcripts (i.e., intron spliced out), the preset used for producing non-spliced, genomic alignments is employed. However, minimap2 parameters could be further optimized for non-spliced alignments to transcriptomes, which have different features than genomes.

StringTie2 and TranSigner were further evaluated on PacBio IsoSeq reads. We used the minimap2-recommended setting for aligning PacBio reads to a genome for benchmarking Bambu and StringTie2 (see Supplementary Text Table 2). As in transcriptomic alignments, we observed that the minimap2 preset for the alignment of PacBio genomic reads caused many of them to be unmapped. In general, TranSigner takes an approach that maximizes the size of the true positive class, as illustrated through the high -N parameter, before EM. For that reason, we decided to keep the ONT genomic alignment preset as it yields more aligned reads. The quality of alignment is critical to the success during TranSigner's EM stage, so further investigation on the optimal minimap2 parameters for transcriptomic alignments is called for.

| | |
|---|---|
| ONT direct RNA | |
| StringTie2, Bambu | `minimap2 -ax splice -uf -k14` |
| NanoCount | `minimap2 -ax map-ont -N 100` |
| TranSigner** | `minimap2 -ax map-ont -N 181` |
| ONT cDNA | |
| StringTie2, Bambu | `minimap2 -ax splice` |
| PacBio IsoSeq | |
| StringTie2 | `minimap2 -ax splice:hq` |

**Supplementary Text Table 2. minimap2 parameters for genomic and transcriptomic alignments**

**TranSigner reuses the same minimap2 parameters for the transcriptomic alignment of all types of long reads.

**3. Short-read-based quantification commands.** As described in Methods, we obtained short-read-based transcript abundance estimates on long-read-derived StringTie2 transcriptomes using Salmon, given paired short and long-read sequencing data on the same biological sample. This analysis involved first running minimap2 to generate spliced genomic alignment of long reads. Different parameters were used for different read types (see Supplementary Text Table 2). Then, StringTie2 was used to obtain a long read-based transcriptome by running:

```
stringtie -L alignments.bam -o transcripts.gtf
```

Next, a Salmon index was created for this StringTie2-assembled transcriptome by:

```
gffread -w transcriptome.fa -g genome.fa transcripts.gtf
```

```
cat transcriptome.fa genome.fa > gentrome.fa
```

```
salmon index -t gentrome.fa -d decoys.txt -i salmon_index
```

Note that the `decoys.txt` file was prepared as instructed in the Salmon documentation. Finally, a Salmon quantification was run as follows:

```
salmon quant -i salmon_index -l A -1 reads_1.fq -2 reads_2.fq -o
salmon_quant --validateMappings
```

**4. Long-read-based quantification commands.** We benchmarked Bambu, FLAIR, StringTie, NanoCount and TranSigner on simulated and experimental long read data. Note that StringTie2 and Bambu require an alignment file as an input, which we prepared using minimap2 (see Supplementary Text Table 2). Bambu and StringTie2 were run as follows:

```
bambu(reads = alignments.bam, annotations = guide.gtf, genome =
genome.fa, trackReads = TRUE)
```

```
stringtie -L alignments.bam -G guide.gtf -o transcripts.gtf
```

For an unguided StringTie2 assembly, we re-used the command introduced in the previous section. Next, FLAIR was run as follows:

```
flair align -g genome.fa -r reads.fq --output flair.aligned
```

```
flair correct -q flair.aligned.bed -g genome.fa -f guide.gtf --output
flair [--nvrna]
```

```
flair collapse -q flair_all_corrected.bed -g genome.fa -r reads.fq --
output flair.collapse --gtf guide.gtf
```

```
flair quantify -r reads_manifest.tsv -i flair.collapse.isoforms.fa --
output flair.quantify --generate_map
```

Note that `--nvrna` flag was added when processing ONT direct RNA reads with FLAIR. Finally, NanoCount was run on the minimap2-generated transcriptome alignments (see Supplementary Text Table 2) as follows:

```
nanocount [-n -d -1] -i alignments.bam -b sel_reads.bam --extra_tx_info
-o tx_counts.tsv
```

Note that `-n -d -1` flags were added when processing ONT cDNA reads with NanoCount, as specified in the tool documentation.

**5. ONT RNA-seq reads simulation.** We simulated ONT direct RNA and cDNA reads using NanoSim. We began the simulation by training a read profile using the NA12878 direct RNA and cDNA reads from Workman et al. using NanoSim's read characterization module as follows:

```
read_analysis.py transcriptome -i reads.fq -rg genome.fa -rt
reference.fa -annot reference.gtf -o out/na12878
```

We used the protein-coding and long non-coding transcripts in the RefSeq v110 annotation of GRCh38 to extract reference.fa and reference.gtf files. NanoSim also requires a quantification TSV file defining the abundances of transcripts to be expressed in the read set. We used Salmon, in its alignment-based mode, to obtain quantify transcript abundances in the Workman et al. NA12878 direct RNA and cDNA reads.

```
minimap2 -ax map-ont reference.fa reads.fq | samtools view -bS >
alignments.bam
```

```
salmon quant -t reference.fa -l A -a alignments.bam -o salmon_quant --
noErrorModel
```

We attempted to run Salmon in its `--ont` mode, but there were errors encountered so we used the `--noErrorModel` instead. We then extracted TPMs from the Salmon output to provide them for read simulation as follows:

```
simulator.py transcriptome -rt reference.fa -rg genome.fa -e
salmon_tpms.tsv -c out/na12878 -o reads -n [14971421, 25418307] -r
[dRNA, cDNA_1D2] --fastq
```

The number of reads specified (i.e., -n flag) corresponds to the number of reads in the Workman et al., NA12878 direct RNA and cDNA samples.

### Supplementary Figures

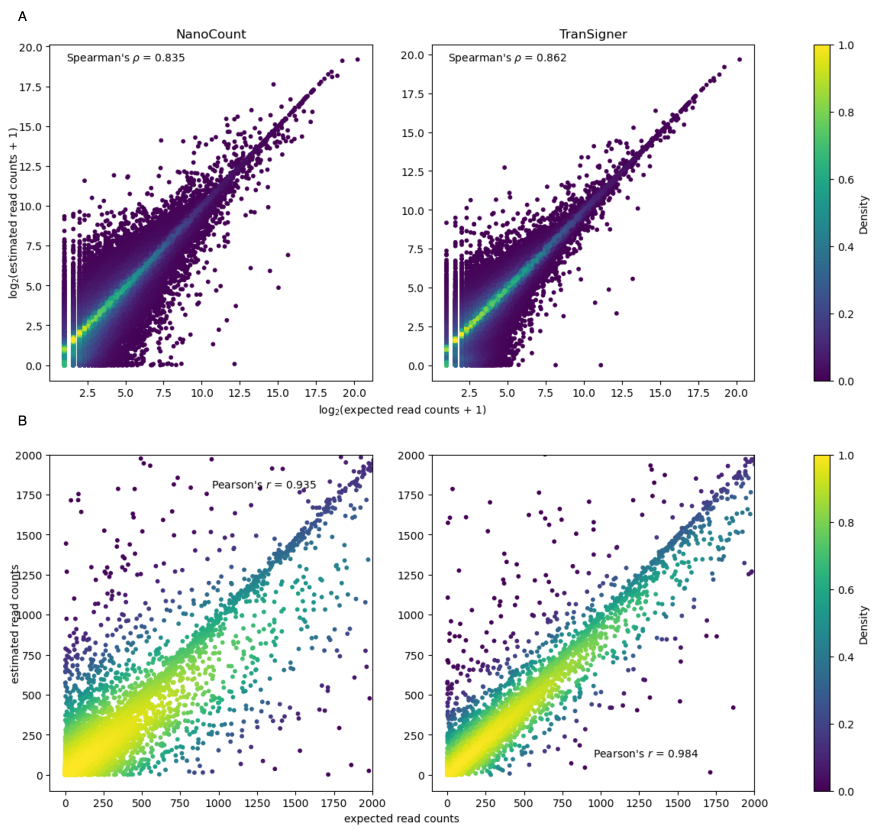


Supplementary Figure S1. Correlation scatter plots comparing ground truth read counts to the read count estimates generated by NanoCount and TranSigner on a simulated ONT cDNA reads set. All tools were provided with the full RefSeq annotation from which the reads were simulated from. A: scatter plots showing the nonlinear correlations between log-transformed expected and estimated read counts. B: scatter plots showing the linear correlations between raw expected and estimated read counts. x- and y-axes were limited to [0, 2000] for demonstration.

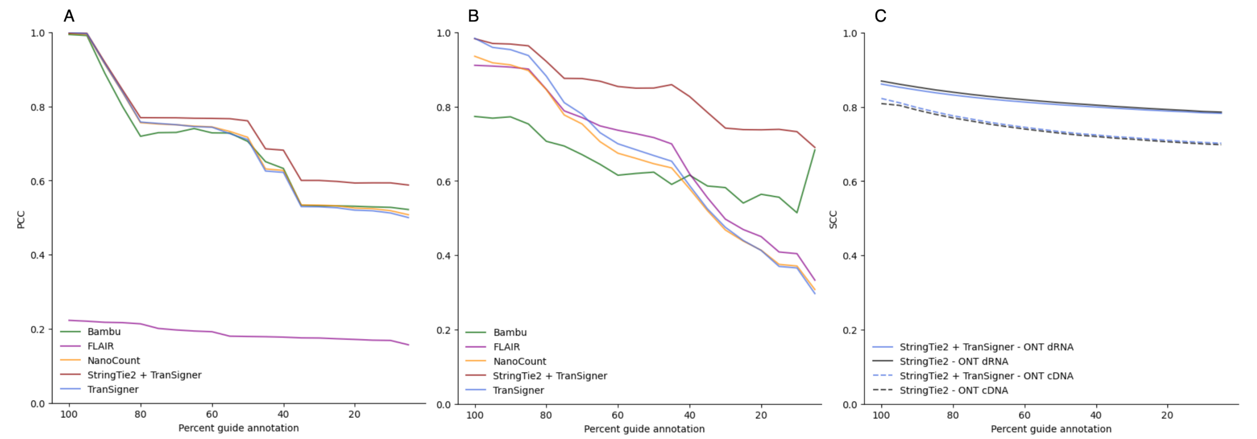


Supplementary Figure S2. Correlation coefficients between true and estimates abundances (read counts in A and B, and per base read coverages in C) computed at varying percent guide annotations using simulated ONT data. A: PCC values in simulated ONT dRNA data. Averages across 9 independent observations (3 read sets, 3 guide samples) shown. B: PCC values in simulated ONT cDNA data. Averages across 6 independent observations (2 read sets, 3 guide samples) shown. C: SCC values for both ONT cDNA (solid line) and cDNA (dotted line) simulated reads. Averages across multiple samples are shown. Different colors indicate different tools.

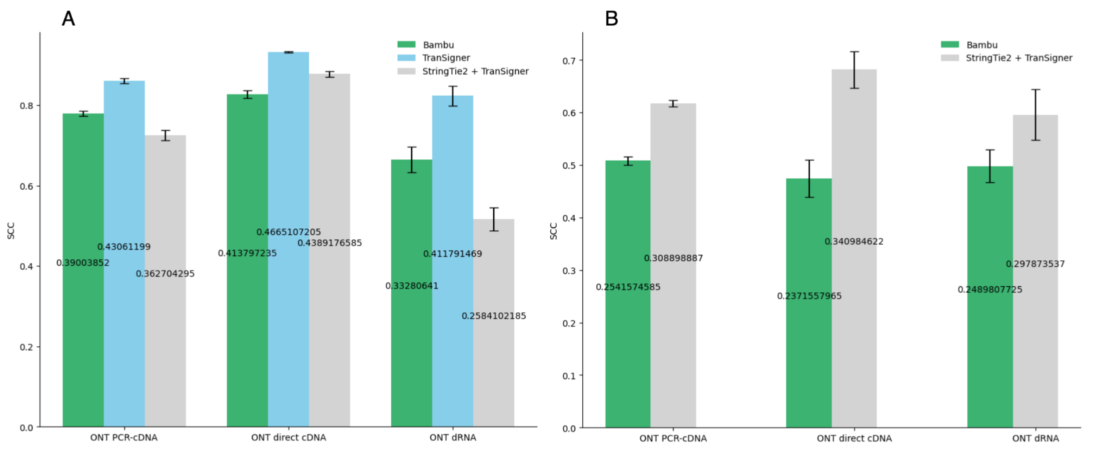


Supplementary Figure S3. Bar plots showing the average SCC values between the expected and estimated sequin CPMs across samples of the same data types. SCCs achieved by Bambu, TranSigner, and StringTie2 A: per read type average SCC values when the full sequin annotation is provided. B: per read type average SCC values when no sequin annotation is provided. Note that TranSigner is missing as it can’t assemble transcripts. Different colors indicates different tools, and error bars show the standard error of measurement (SEM) values.

**ALGORITHM 1: DROP**

**Input**: a compatibility scores matrix $X$ and a matrix containing read fractions $\alpha$
**Output**: updated compatibility scores matrix $X$

for $r$ = 1, … , $N$ do (iterate through all reads in $\alpha$)
    $M \leftarrow$ length($\alpha_r$)    // $|\alpha_r|$ equal to $|T_r|$ in Methods
    $k \leftarrow 1 / M$    // $k$ equal to $\tau_r$ in Methods
    for $t$ = 1, … , $M$ do (iterate through all transcripts aligned to a read)
        if $\alpha_{rt} > k$ then
            $X_{rt} \leftarrow 0$
        else
            do nothing
        end
    end
end

Supplementary Figure S4. Pseudocode for TranSigner's drop algorithm. The input to this algorithm is two matrics, one storing the compatibility scores between reads and transcripts ($X$) and another containing the read fractions assigned to transcripts ($\alpha$). The output of this algorithm is the updated compatibility scores matrix $X$ that gets used to perform all subsequent E-steps, which isn't shown above but described in Methods. A drop occurs when the fraction of a read $r$ assigned to a transcript $t$ is less than some threshold. It removes the compatibility relationship between a read $r$ and a transcript $t$ is removed by setting the $X_{rt}$ to 0 to ensure that $\alpha_{rt}$ remains 0 (i.e., no fraction of $r$ assigned to $t$) in subsequent E-step updates as it's a product of $X_{rt}$. The drop threshold is dynamically computed for each read $r$ as 1 / $M$ where $M$ is the number of transcripts compatible with $r$.

**ALGORITHM 2: PUSH**

**Input**: a matrix storing the read fractions $\alpha$, and relative transcript abundances $\rho$
**Output**: a list $\alpha'$ containing hard assignments between reads and transcripts, new set of relative transcript abundances $\rho'$ computed using those hard assignments

$\alpha' \leftarrow$ an empty list of size $N$ (initialize $\alpha'$)
for $i = 1, \ldots, N$ do (iterate through all reads)
  $M \leftarrow \text{length}(\alpha_i)$
  $j \leftarrow \underset{j}{\text{argmax}}\, \alpha_{ij}$
  $\alpha'[i] \leftarrow j$
end
// perform M-step updates to obtain the updated relative transcript abundances
$\rho' \leftarrow \text{M\_step}(\alpha')$

Supplementary Figure S5. Pseudocode for TranSigner's push algorithm that can be used to obtain 1-to-1 hard assignments between reads and transcripts. The input to this algorithm is a matrix containing read fractions assigned to transcripts ($\alpha$) and the relative transcirpt abundances ($\rho$). The output of this algorithm is a list containing hard read-to-transcript assignments ($\alpha'$) and new relative abundances computed using these new assignments ($\rho'$). The push algorithm assigns each read $r$ to the transcript with the most read fraction assigned, among all transcripts compatible with it.
